# Redefining Genomic Privacy: Trust and Empowerment

**DOI:** 10.1101/006601

**Authors:** Arvind Narayana, Kenneth Yocum, David Glazer, Nita Farahany, Maynard Olson, Lincoln D. Stein, James B. Williams, Jan A. Witkowski, Robert C. Kain, Yaniv Erlich

**Affiliations:** Department of Computer Science, Princeton University, 35 Olden Street, Princeton, New Jersey 08540, USA; Illumina Inc. 5200 Illumina Way, San Diego, California 92121, USA; Google Inc, 1600 Amphitheatre Parkway, Mountain View, California, 94043, USA; Duke University School of Law; Duke Science & Society, 210 Science Drive, Box 90362, Durham NC 27705, USA; University of Washington, P.O. Box 339 Port Orford, Oregon 97465, USA; Ontario Institute for Cancer Research, 661 University Avenue, Suite 510 Toronto, Ontario Canada, M5G 0A3; Department of Molecular Genetics, University of Toronto, 100 St. George Street, Toronto, Ontario Canada M5S 3G3; Banbury Center, Cold Spring Harbor Laboratory, One Bungtown Road NY 11724, USA; Whitehead Institute for Biomedical Research, Nine Cambridge Center, Cambridge, MA 02142, USA

## Abstract

Fulfilling the promise of the genetic revolution requires the analysis of large datasets containing information from thousands to millions of participants. However, sharing human genomic data requires protecting subjects from potential harm. Current models rely on de-identification techniques that treat privacy versus data utility as a zero-sum game. Instead we propose using trust-enabling techniques to create a solution where researchers and participants both win. To do so we introduce three principles that facilitate trust in genetic research and outline one possible framework built upon those principles. Our hope is that such trust-centric frameworks provide a sustainable solution that reconciles genetic privacy with data sharing and facilitates genetic research.

## 1 Introduction

> ”Widespread distrust … imposes a kind of tax on all forms of economic activity, a tax that high-trust societies do not have to pay”
>
> — — *Francis Fukuyama* [1]

Genomic research promises substantial societal benefits, improving health care and our understanding of human biology, behavior, and history. To deliver on this promise, the research and medical community require active participation of a large number of human volunteers and broad dissemination of genetic datasets. However there are serious concerns about potential abuses of genomic information, from racial discrimination and denial of services due to genetic predispositions, to disclosure of intimate familial relationships such as non-paternity events. Todays data-management techniques largely frame the value of data versus the risks to participants as a zero-sum game, in which one players gain is anothers loss.

Facing these challenges, we recently held a meeting at the Banbury Center on Accelerating Genomic Research with Privacy Protections (Dec 11-12, 2013). The aim of the meeting was to discuss policy and technical improvements and alternatives to the status quo. The meeting brought together experts in computer science, data privacy, bioethics, policy, law enforcement, consumer rights, bioinformatics, and human genetics. This manuscript distills major points from the meeting and extensive follow-on discussions between the participants.

## 2 The rise and fall of de-identification

Current models for protecting participant data in genetic studies focus on concealing the participants identities. This focus is codified in the legal and ethical frameworks that govern research activities in most countries. Most data-protection regimes were designed to allow the free flow of de-identified data, while restricting the flow of personal information. For instance, both the US HIPAA rule and the EU Privacy directive require either explicit subject consent or proof of minimized risk of re-identification before data dissemination. In Canada, the test for whether there is a risk of identification involves ascertaining whether there is a “serious possibility that an individual could be identified through the use of that information, alone or in combination with other available information.” [2] To that end, the research community employs a fragmented system to enforce privacy that includes *institutional review boards* (IRB), ad-hoc *data access committees* (DAC), and a range of privacy and security practices such as HIPAA Safe Harbor.

However, the current protection-by-anonymity paradigm is under threat. Standard data security controls are important but not sufficient for genetic data. For instance, access control and encryption can ensure the security of information at rest in the same fashion as for other sensitive (e.g. financial) information, protecting against outsiders or unauthorized users gaining access to data. However, there is also a need to prevent misuse of data by a ‘legitimate’ data recipient. Recent advances in re-identification attacks, specifically against genetic information, reduce the utility of these de-identification techniques [3].

With the growing limitations to de-identification, the current paradigm is not sustainable. At best, participants go through a lengthy, cumbersome, and poorly understood consent process that tries to predict worst-case future harm. At worst, they receive broken promises for anonymity. Data custodians must keep maneuvering between the opposite demands for data utility and privacy, relegating genetic datasets into silos with arbitrary access rules. Funding agencies waste resources funding studies whose datasets cannot be reused across and between large patient communities because of security concerns. Finally, well-intentioned researchers struggle to obtain genetic data from hard to access resources. These limitations impede serendipitous and innovative research and degrade a data set’s research value, with published results often overturned due to small sample sizes [4].

### The gaps in current data privacy techniques

It may be that current de-identification practice, which primarily consists of removing an individuals personally identifying information from records containing individualized genetic information, is simply outdated; though a war of attrition, it is possible that new techniques will once more make it difficult to re-identify an individual. In the meeting, we reviewed computational schemes that theoretically make re-identification demonstrably (and perhaps quantifiably) difficult. We focused on two classes of techniques: *secure multiparty computation* (MPC) [5] – including *homomorphic encryption* – and *differential privacy* (DP) [6]. The hallmark of these techniques is that they provide mathematical proofs delineating what the data recipient can and cannot infer based on the data access given to them.

MPC computes a known, shared function on encrypted data sets from multiple parties; the computation reveals nothing about the parties input data other than the functions results. For example, a patient or her physician holding genetic data can use MPC to have their genetic data interpreted by a third party service without revealing the actual genotypes. However, MPC has some practical limitations. First of all, MPC requires predefined analysis protocols. Unfortunately, research protocols are rarely fixed in advance. Most research is exploratory in nature, and is characterized by ad-hoc analyses in which researchers test and refine their analytic procedures repeatedly during the course of the study. Moreover, MPC does not address privacy breaches such as attribute-disclosure attacks (e.g., the Homer et al. study [7]) that result from the output of the analytic procedure rather than its inputs.

In contrast, differential privacy (DP) applies to the scenario of a trusted party holding a database of sensitive information wishing to publish the result of a function computed on the data. Techniques used to achieve DP include the addition of noise to the output. Researchers have developed DP algorithms for performing many data-mining tasks, such as reporting summary statistics and clustering. Unfortunately, the current levels of noise required for differential privacy appear to be unacceptable for most genetic studies and would eradicate the weak association signals that are the reality of most complex traits.

Our conclusion is that these emerging computational techniques for ensuring genetic privacy show potential, but require substantial theoretical and practical development to be fully operational methods for data sharing to accelerate scientific studies.

## 3 Focusing on trust not privacy

We propose to shift from the zero-sum game of data privacy versus data utility to a framework that builds and maintains *trust* between participants and researchers. We suggest the following key principles for trust-enabling frameworks:

1. **Transparency creates trust**: Trust requires transparency between parties. In genomic research, transparency means informing participants about not only the intended but the actual use of data. This is a commonly accepted principle of information privacy that is found in most data protection statutes (e.g., Canadas PIPEDA [8]) and fair information practices (e.g., the OECD Privacy Principles [9]).
2. **Increased control enhances trust**: Given the uncertainties in genetic studies, the burden of making “fully informed” decisions about future data use and harms is virtually impossible. However, the situation improves when the participant is given control over future data use. Clear communication of risks is crucial to ensure fully-informed participants, yet current consent processes require participants to make a one-time decision about future data sharing preferences with unknown risks. Even worse, some consent forms include vague ‘legalese’ that might be tempting from a legal perspective, but instead fuel patients fears. Some participants naturally shy away from sharing when the terms are too broad, while other individuals might make decisions that are not well informed. In addition, one-time blanket consent does not accommodate the reality that privacy preferences might change over time.
3. **Reciprocity maintains trust**: Researchers should maximize the value of data collected from participants, subject to individual preferences. By advancing scientific knowledge, the research community reciprocates and pays back the participants volunteerism. A sense of community among participants can help bridge the gap between societal and individual rewards. Mechanisms for participants to reward researches that act appropriately and punish researchers that violate their trust provide incentives for ongoing win-win behavior.

If successful, a trust-centric framework creates a system that rewards good behavior, deters malicious behavior, and punishes non-compliance. This stands in stark contrast to the current system that punishes researchers, participants, and progress.

## 4 Bilateral Consent Framework

Building on top of the three key principles above, we suggest a trust-enabling framework, called the *Bilateral Consent Framework* (BCF) (Table 1). This approach is inspired by the recent movement for participant-centered research [10] and the growing success of trust-enabling techniques in online peer-to-peer economy marketplaces. Importantly, our proposal is not meant to be final, but rather to provide a framework and a set of building blocks to drive discussions among the community. The major building blocks of the BCF are the following:

**Table 1:** Major differences between current data sharing frameworks and a BCF.

| Attribute | Current system | BCF |
| --- | --- | --- |
| Consent for secondary use | One time decision | Dynamic |
| Primary data controller | PI | Participant |
| Who decides on secondary data usage? | Data Access Committee or local IRB | Participant |
| Data stewardship | Not defined | Trusted mediator |
| Code of conduct | Locally determined | Globally determined |
| Oversight | Local IRB | The community (participants, trusted mediator, researchers) |
| Oversight mechanism | Not clear | Audit system |
| Who can punish data misconduct? | Local IRB | The community (participants, trusted mediator, researchers) |
| Main source of reputation is | University or research institute | The community (previous participants, trusted mediator, researchers) |
| Cohort integrity | Stable | Indefinite / variable |
| Place of computation | PI-owned equipment or PI-chosen cloud provider | Resource-owned equipment or resource-chosen cloud provider. |

*Trusted mediator*: the role of the trusted mediator is to operate the BCF. This entity can be a patient-advocacy group, a funding agency, academic center, a scientific society, or a private company as long as they are trusted by the participants and have the means to operate the BCF. The trusted entity should mediate the communication between the researchers and the participants, act upon the participants decisions, and be the single point of contact. In addition, this entity should educate participants about the nature of the data and describe the benefits and risks.

*Dynamic participant consent*: at its core BCF enables participants to have enhanced and dynamic control over access to data about them. In current consent architectures, the participant delegates complete control over the data to the Principal Investigators (PIs). Upon completion of the study, the PI typically delegates secondary usage decisions to a data access committee or an IRB board. In the BCF, data control remains primarily tied to the source individual. Researchers solicit their studies, describing the benefits of the study and specifying limitations on how they use the data. The participant can grant or deny consent to different studies. Thus, instead of one-time decisions about data sharing, a BCF fosters long-term engagement by participants, allowing researchers to solicit participant data and participants to change their data contribution as they see fit.

*Uniform code of conduct*: having researchers consent to uniform guidelines makes it easier for participants to grant consent to new researchers. Researchers that are part of the BCF consent to a code of conduct that affirms that individual data will be properly handled, including that it will be held securely and that re-identification will not be attempted. Thus, BCF replaces the gatekeeer approach, where IRBs decide who should count as a qualified researcher on a case-by-case basis, with a participant-centric model, where participants understand the rules that researchers will follow. Evidence for violation of the code of conduct can result in public notice, canceled access, and possible legal action. Methods for redress might include data protection law, criminal law or additional contractual terms (e.g. indemnification and compensation).

*Auditing*: the BCF encourages a trust-but-verify approach. All data access should be monitored, both to remind researchers that their access privileges depend on trust, and to enable potential detection and enforcement of violations. One means of monitoring is for all analysis activity to be executed on the trusted mediators computing resources and logged. This is different from current access control models where upon permission the researcher analyzes the data on his or her own computing resources without any oversight on the actual analysis. Importantly, we do not expect the auditing system to be perfect or to capture all data misuse. The primary aim of such system is to deter malicious behavior. However, we envision that in the future such systems can help to identify clear anomalies (e.g. analysis of Y-STRs that is a key component of surname inference [11]) or data analysis that is substantially different from the consent. In addition, logging and auditing promote transparency. There is growing interest in using cloud computing for genetic analysis and moving the computation to the data; adding an auditing system can leverage this trend to increase trust.

*Reputation system*: having information about researchers past behavior will help participants make good consent decisions. We propose a reputation system that rewards researchers that maintain solid records of adhering to their promise for how data entrusted to their care is used. Such systems have catalyzed online marketplaces that require high levels of trust with minimal previous interaction, such as Ebay or Uber. In the BCF, a reputation system can include the researchers portfolio of previously completed studies, recommendations from previous participants, and a vouching system from other researchers. Accordingly, participants can elect to share data only with researchers of sufficient reputation and the trusted entity can revoke access to researchers with low reputation.

The aim of the description above is to describe potential architectural elements of trust-centric frameworks. While these building blocks reinforce each other, they are not meant to be an all-or-nothing monolithic system. We recognize that certain implementations might only use a subset of these elements.

## 5 Conclusion

Realizing a bilateral consent framework will require new technologies and hard choices. However, there is a need for improved global standards for legal and technical frameworks to share genomic data. Initiatives such as the Global Alliance for Genomics and Health [12] and the Genetic Alliance [13] have started the dialogue; it is our hope that the proposed framework can act as a starting point as stakeholders move from discussion to practice. A bilateral consent framework can redirect fears of unknown privacy abuse to excitement for participating in the genetic information revolution.

## Acknowledgments

The authors thank the Banbury Center for hosting the meeting, Illumina Inc. for the funding that enabled the meeting, and Steven Brenner and Laura Rodriguez for useful comments. YE, RK, AN, and JAW organized the meeting. LDS is the recipient of awards from Ontario Institute for Cancer Research, the National Institutes of Health, the Canadian Institutes of Health Research, the National Scientific and Engineering Research Council of Canada, the Ontario Research Fund, and Genome Canada. YE is an Andria and Paul Heafy Family Fellow and holds a Career Award at the Scientific Interface from the Burroughs Wellcome Fund.

